# Rats remember: Lack of drug-induced post-retrieval amnesia for auditory fear memories

**DOI:** 10.1101/2020.07.08.193383

**Authors:** Laura Luyten, Anna Elisabeth Schnell, Natalie Schroyens, Tom Beckers

## Abstract

When retrieved under specific circumstances, consolidated fear memories are thought to return to a labile state, thereby opening a window for modification (e.g., attenuation) of the memory. Several interventions during a critical time frame after this destabilization seem to be able to alter the retrieved memory, for example through pharmacological interference with the restabilization process, either by direct protein synthesis inhibition or indirectly, using drugs that can be safely administered in patients (e.g., propranolol).

In a series of well-powered auditory fear conditioning experiments (four with propranolol, 10 mg/kg, two with rapamycin, 20-40 mg/kg, one with anisomycin, 150 mg/kg and cycloheximide, 1.5 mg/kg), we found no evidence for reduced cued fear memory expression during a drug-free test in adult male Sprague-Dawley rats that had previously received a systemic drug injection upon retrieval of the tone fear memory. All experiments used standard fear conditioning and reactivation procedures with freezing as the behavioral read-out (conceptual or exact replications of published reports) and common pharmacological agents. Additional tests confirmed that the applied drug doses and administration routes were effective in inducing their conventional effects on expression of fear (propranolol, acutely), body weight (rapamycin, anisomycin, cycloheximide) and consolidation of extinction memories (cycloheximide).

Thus, in contrast with most published studies, we did not find evidence for drug-induced post-retrieval amnesia, underlining that this effect, as well as its clinical applicability, may be considerably more constrained and less readily reproduced than what the current literature would suggest.

**Highlights:**

- We aimed to replicate post-retrieval amnesia for auditory fear memories in rats
- We performed a series of well-powered pharmacological interference experiments
- Propranolol, rapamycin, anisomycin or cycloheximide was injected upon retrieval
- Bayesian stats found substantial evidence for the absence of post-retrieval amnesia
- The effect is less reproducible and generalizable than what the literature suggests

## Introduction

Consolidated memories have long been seen as immutable, but accumulating evidence suggests that they are not set in stone and that they can still be modified or even erased after the completion of synaptic consolidation (Dudai, 2004; McGaugh, 2000). This paradigm shift was first proposed in the late 1960’s (Misanin, Miller, & Lewis, 1968) and received important support 20 years ago (Nader, Schafe, & LeDoux, 2000; Przybyslawski, Roullet, & Sara, 1999). These studies suggested that interventions such as an electroconvulsive shock or drug administration into the basolateral amygdala shortly after reactivation of the memory allowed for interference with this memory in such a way that there was a significant attenuation of memory expression (i.e., amnesia) on subsequent testing. It is not within the scope of this paper to give a comprehensive overview of all these findings (for a review, see e.g., Beckers & Kindt, 2017), but it is safe to say that many of these studies have put forward that this technique could be a game changer in the treatment of several types of psychopathology in which maladaptive memories are a core feature, e.g., post-traumatic stress disorder (PTSD), other anxiety-related disorders, and even substance use disorders (Beckers & Kindt, 2017; Przybyslawski, et al., 1999). Many studies have examined the neural substrate of the observed post-retrieval amnesia through local manipulations in specific brain areas (e.g., Debiec, LeDoux, & Nader, 2002; Nader, et al., 2000), while others have relied on the use of systemic pharmacological manipulations following memory reactivation (e.g., Debiec & LeDoux, 2004; Gisquet-Verrier et al., 2015; Tallot et al., 2017; Taubenfeld, Milekic, Monti, & Alberini, 2001). Whereas local administration obviously provides higher spatial accuracy, disadvantages may include unwanted effects of protein synthesis inhibitors, such as cell death at the site of injection, and the need for specific assumptions regarding the key region of interest (Flint, Valentine, & Papandrea, 2007). An obvious advantage of systemic pharmacological manipulations, on the other hand, is the higher translatability of findings to clinical applications. A prime example is the use of propranolol for post-retrieval attenuation of fear memories, first shown in fear-conditioned rodents (Debiec & LeDoux, 2004), and later also in fear-conditioned humans (Kindt, Soeter, & Vervliet, 2009, but see Bos, Beckers, & Kindt, 2014) and spider phobics (Soeter & Kindt, 2015). Propranolol is a centrally acting beta-adrenergic antagonist and has repeatedly been shown to be effective for the induction of amnesia upon reactivation of the targeted memory. Although safe and apparently successful, propranolol seems to be the odd one out in light of the widely supported (but sometimes contested, Gisquet-Verrier & Riccio, 2018) mechanism underlying the observed amnesia, i.e., reconsolidation interference. The reconsolidation hypothesis states that a consolidated memory can re-enter a labile phase through reactivation, which then requires protein synthesis (during reconsolidation) in order to preserve the original memory (Nader, et al., 2000). Given the proposed prerequisite of protein synthesis, the most obvious way to interfere with such reconsolidation is through administration of protein synthesis inhibitors, which have been used often and with success, although their profile is much more toxic than that of propranolol (which is assumed to have indirect effects on protein synthesis (Przybyslawski, et al., 1999)). Commonly used protein synthesis inhibitors in this field of research include anisomycin, cycloheximide and rapamycin. Systemic anisomycin, for instance, has been successfully used for induction of post-retrieval amnesia in several behavioral procedures (e.g., context conditioning, conditioned place preference) in numerous rodent studies (Bernardi, et al., 2007; Blundell, Kouser, & Powell, 2008; Fan et al., 2010; Lattal & Abel, 2004; Milekic, Brown, Castellini, & Alberini, 2006; Suzuki, et al., 2004 are only a few examples).

A few years ago, we set out to optimize a protocol to investigate post-retrieval amnesia in rat fear conditioning in our laboratory. Auditory fear conditioning is a germane tool to study mechanisms central to anxiety-related disorders, such as PTSD and phobias, and was therefore the focus of our effort. The literature seems to suggest that drug-induced post-retrieval amnesia is relatively easy to obtain, considering the many successful studies with rats and mice, using a plethora of different pharmacological agents (Beckers & Kindt, 2017; Reichelt & Lee, 2013). Furthermore, to our knowledge, there are no published failures to find such a drug-induced amnestic effect with rodent auditory fear conditioning. Several studies do indicate that the effect is not, or less easily, found with old or strong memories (e.g., Wang, de Oliveira Alvares, & Nader, 2009 (auditory fear), Milekic & Alberini, 2002 (inhibitory avoidance), Bustos, Maldonado, & Molina, 2009; Suzuki et al., 2004 (contextual fear)), but all these papers do find evidence for amnesia under ‘standard conditions’, i.e., recent fear memories that are reactivated through a typical protocol (e.g., one unreinforced presentation of the conditioned stimulus), after which administration of the drug results in a considerable deficit in memory expression, usually tested one day later (Tronson & Taylor, 2007).

Of note, published studies using systemic or intra-amygdala injection of propranolol, anisomycin or cycloheximide typically report (very) large effects (e.g., Debiec & LeDoux, 2004; Duvarci, Nader, & LeDoux, 2005; Gisquet-Verrier et al., 2015; Muravieva & Alberini, 2010; Nader, et al., 2000; Taubenfeld, et al., 2001). Considering these effect sizes, the studies presented here are well-powered to detect differences between drug-treated and control groups. Published effect sizes with rapamycin are considerably smaller (e.g., Hoffman et al., 2015; Y. Li et al., 2013; Tallot, et al., 2017), but this protein synthesis inhibitor has the benefit of typically being administered systemically (see also e.g., Barak et al., 2013; Blundell, Kouser, & Powell, 2008; Mac Callum, Hebert, Adamec, & Blundell, 2014). Moreover, rapamycin and analogs are approved for use in transplant and cancer patients, making it a worthwhile candidate to explore in view of future clinical applications (J. Li, Kim, & Blenis, 2014).

To summarize, we conducted a series of sufficiently powered auditory fear conditioning experiments in adult male rats (see ‘Power calculations’ for details), in which we aimed to induce amnesia by systemically administering one of four drugs (propranolol, rapamycin, anisomycin or cycloheximide) after retrieval of the tone fear memory. Experimental parameters (e.g., shock intensity, testing conditions) were varied slightly in between experiments in order to optimize our chances of finding the effect, and included an exact replication of Debiec & LeDoux (2004). In addition, we carried out control experiments and analyses to confirm that the drugs were biologically active at the applied dose and administration route. More specifically, as a positive control, we investigated the effect of cycloheximide on the consolidation of fear memories, by administering the drug immediately after training, rather than after reactivation, in line with published studies which again report large effect sizes (Kochli, Thompson, Fricke, Postle, & Quinn, 2015; Lay, Westbrook, Glanzman, & Holmes, 2018; Stiedl, Palve, Radulovic, Birkenfeld, & Spiess, 1999). In a subset of animals, we also evaluated the effect of cycloheximide on the consolidation of an extinction memory.

## Materials & Methods

### Preregistration and data availability

All experiments, including protocols, planned sample sizes and analysis plans, were registered on the Open Science Framework (OSF) before the start of data collection (https://osf.io/j5dgx). Raw data can be found there as well.

### Subjects

One hundred sixty-four male Sprague-Dawley rats (270-300 g on arrival in the lab) (Janvier Labs, Le Genest-Saint-Isle, France) were housed individually in plastic cages with bedding, food and water ad libitum. This project was approved by the animal ethics committee at KU Leuven and is in accordance with the Belgian and European laws, guidelines and policies for animal experimentation, housing and care (Belgian Royal Decree of 29 May 2013 and European Directive 2010/63/EU on the protection of animals used for scientific purposes of 20 October 2010). Cage enrichment (Dura-Chew, Bio-Serv, Flemington, NJ, USA) was provided in Experiments 2-5. To monitor general wellbeing, animals were weighed at several points before, during and/or after the experiments, but never on Training (except for Experiment 1), Reactivation or Test days, in order to disturb the behavioral sessions as little as possible. Body weight on the Reactivation day (for calculation of the drug injection volume) was estimated as the most recent weight + 10 g/day. Experiments started 3-5 days after arrival in the lab. Animals were kept on a 12h/12h day-night cycle, with experiments starting 1-2.5 hours after the beginning of the light phase. Rats were brought from the housing facility to the testing room approximately 2 minutes before each behavioral session in their home cage, carried by the experimenter (Experiment 1) or transported on a wheeled cart (all other experiments).

### Procedure

Experimental designs, including the number of tones (conditioned stimulus, CS) and shocks (unconditioned stimulus, US) presented during each session, are shown in ***Fig. 1-3***. Specific details regarding tone and shock parameters and timing of the stimuli can be found in the ***Supplement***. In all experiments, freezing was scored manually from videos by an experienced observer blinded to group allocation (Luyten, Schroyens, Hermans, & Beckers, 2014), and expressed as a percentage of the time under evaluation. Preregistered criteria for exclusion were less than 15% freezing during the Reactivation CS in Experiments 4 and 7, and exclusion from Test 2 analyses in Experiment 7 if freezing was less than 15% during the first 3 CSs of Test 1.

**Figure 1:**
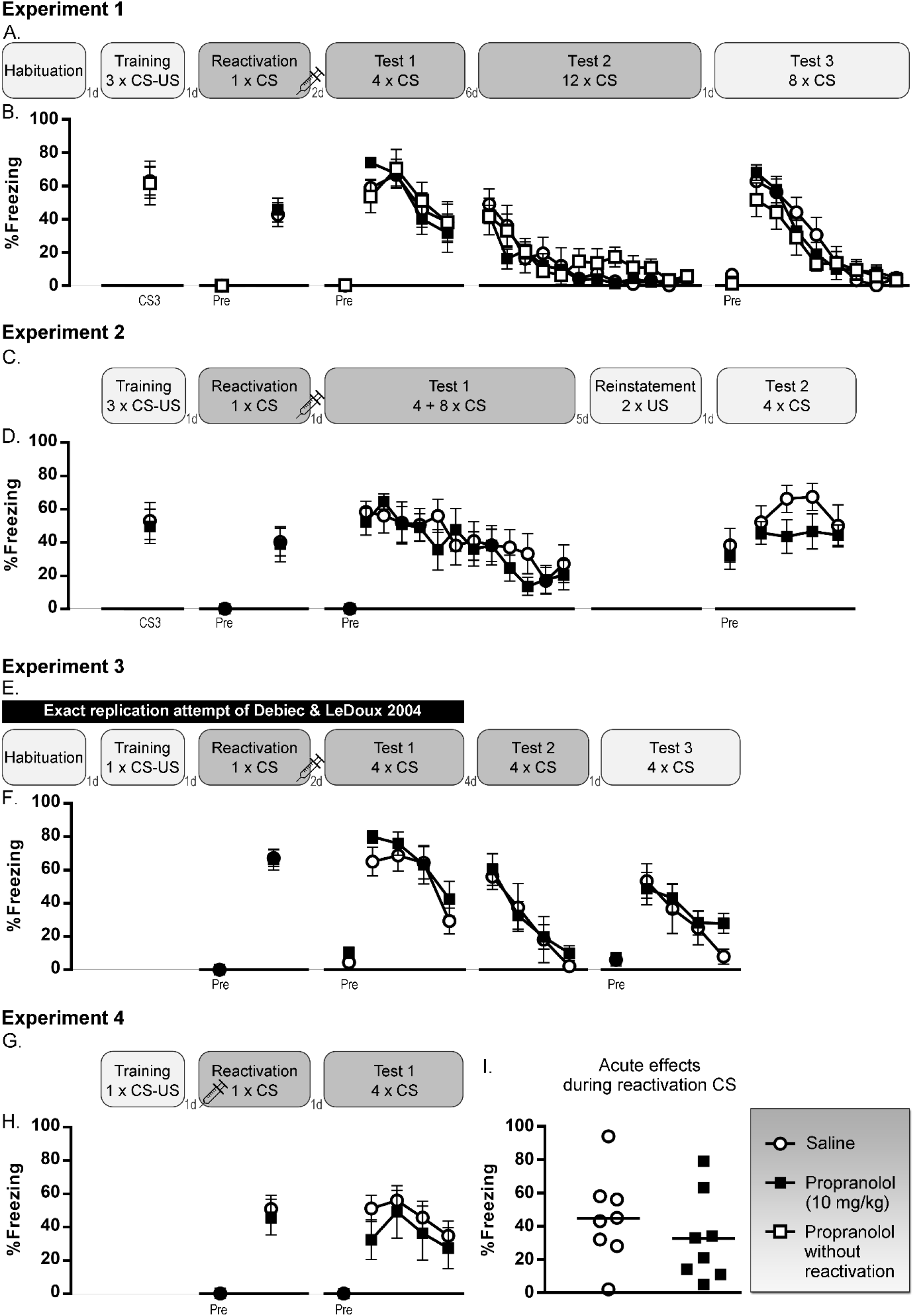
Propranolol (10 mg/kg) experiments. **A-B.** Experiment 1, n = 8 per group. **C-D.** Experiment 2, n = 8 per group. **E-F.** Experiment 3, n = 6 saline rats and n = 8 propranolol rats. **G-H.** Experiment 4, n = 7 saline rats and n = 5 propranolol rats. Percentage freezing during each tone (mean and SEM) is shown. ‘Pre’ is contextual freezing before the first tone presentation of the session. **I.** Acute effects of propranolol in Experiment 4, in which the drug was administered immediately before rather than after reactivation, n = 8 per group. Individual data and group means are shown. Light gray box indicates that a session takes place in context A, dark gray is context B. CS: conditioned stimulus, US: unconditioned stimulus, d: day(s).

### Equipment

Four identical chambers (Contextual NIR Video Fear Conditioning System for Rats, Med Associates Inc., St. Albans, VT, USA) were equipped with specific contextual features for Context A and Context B. Built-in ventilation fans provided background noise (±67 dB) in all chambers. Context A had a standard grid floor, a black triangular insert, was illuminated by infrared and white light (50 lux) and cleaned and scented with a pine odor cleaning product. Context B had a white plastic floor, a white plastic curved back wall insert, infrared light only, and was cleaned and scented with a different cleaning product.

### Experiments 1-7

#### Habituation

. In some experiments, animals were habituated to Context A (Experiments 1 and 3), or Context A and B (Experiment 7) one day before Training.

#### Training

To acquire a cued fear memory, rats received 1 or 3 tone-shock pairings in Context A. Tones were 5000 Hz, 75-80 dB for 20-30 s and co-terminated with the shocks (0.7-1.5 mA, 0.5-1 s, depending on the experiment, see ***Supplement***). In experiments with 3 CS-US pairings, animals were matched into equivalent groups (block-randomization) based upon freezing during the third CS.

#### Reactivation

Twenty-four hours later, a single tone CS without shock was presented in Context B with the aim of reactivating the cued fear memory and rendering it vulnerable to drug interference. Approximately 1-2 minutes after the end of the Reactivation session, the drug or its control vehicle was injected systemically. In Experiment 4, the injection was given 3-6 min before (instead of after) Reactivation. Having the propranolol on board during the reactivation phase may, theoretically, increase the opportunities to block reconsolidation already in the earliest stages after reactivation (Kindt, et al., 2009; Muravieva & Alberini, 2010; Vetere et al., 2013). In Experiment 7, short-term memory was tested in Context B, 4 hours after Reactivation (no amnestic effects were expected here).

#### Tests

One or two (in Experiments 1 and 3) days later, retention of the tone fear memory was tested in Context B (Test 1), to evaluate the anticipated drug-induced post-retrieval amnesia. This test consisted of 3 (in Experiment 7) or 4 CS presentations. Note that in Experiments 2, 5 and 6, more CSs were presented with the aim of inducing extinction. Nevertheless, the use of the first 3-4 CSs as an index of long-term fear memory retention was preregistered. Under the assumption that we were going to induce post-retrieval amnesia, most experiments featured additional tests to assess the robustness of the amnesia that we were expecting to see, including spontaneous recovery tests after (partial) extinction of the fear memory in Experiments 1, 3 and 5, renewal tests in context A in Experiments 1, 3, 5 and 7 and reinstatement tests in Experiments 2 and 6.

#### Exact replications

Experiment 3 (up to and including Test 1) was an exact replication of Debiec & LeDoux (2004). In addition, the behavioral procedure (up to Test 1) of Experiment 7 was an exact replication of a protocol used in Duvarci et al. (2005) and Nader et al. (2000) (details see ***Supplement***). The route of drug administration did, however, differ between their studies (infusion into the basolateral amygdala) and ours, as we chose to use systemic injections throughout. All other experiments were conceptual replications of prior successful studies.

### Experiment 8

Experiment 8 was designed to test drug effects on consolidation rather than reconsolidation. Therefore, on the first day, training of a cued fear memory with one tone-shock pairing took place in context A, immediately followed by injection of cycloheximide or vehicle. One day later, retention of the cued fear memory (and possible effects of cycloheximide on its consolidation) was assessed during the first 3 CSs of Test 1 in context B. This test session continued with further extinction of the cued fear memory. Immediately after this extinction training, rats that had received vehicle after Training, now received cycloheximide or vehicle. One day later, retention of the extinction memory (and possible effects of cycloheximide on its consolidation) in this subset was assessed during Test 2 in context B.

### Drugs

#### Propranolol

In Experiments 1-4, propranolol (Product P0884, Sigma-Aldrich, Overijse, Belgium) was dissolved in saline on the day of injection to obtain a solution of 10 mg/ml, administered intraperitoneally at 1 ml/kg (10 mg/kg dose). Control animals received saline (1 ml/kg). Injections were given in the testing room, except for Experiment 4 where animals were injected in an adjacent room.

#### Rapamycin

In Experiments 5-6, rapamycin (LC Laboratories, Woburn, MA, USA) was dissolved in vehicle (100% DMSO) (99.8% dimethyl sulfoxide extra pure, Acros Organics, Geel, Belgium) on the day of injection to obtain a solution of 20 mg/ml, administered intraperitoneally at 1 ml/kg (20 mg/kg dose in Experiment 5) or 2 ml/kg (40 mg/kg dose in Experiment 6). Injections were given in the testing room.

#### Anisomycin & Cycloheximide

In Experiments 7-8, anisomycin (Product A9789, Sigma-Aldrich) (50 mg/ml) was dissolved in saline which was brought to pH ≤5 using HCl, and then adjusted again to pH ±7-7.4 with NaOH. Cycloheximide (Product C7698, Sigma Aldrich) (0.5 mg/ml) and the vehicle solution were prepared using the same procedure. All solutions were made on the day before injection and stored in the fridge until 30 minutes before injection, while continuously being shielded from light. Solutions were administered subcutaneously in the nape of the neck at 3 ml/kg (150 mg/kg dose of anisomycin or 1.5 mg/kg dose of cycloheximide). Injections were given in a room adjacent to the testing room. Note that Gisquet-Verrier et al. (2015) used 2.8 mg/kg cycloheximide intraperitoneally, but mentioned the loss of several animals, which is why we decided to use a lower dose. The doses applied in this study are around the tolerance threshold (see ***Supplement***), and these (or sometimes lower) doses have been shown to be effective in other rat learning procedures (e.g., Bernardi, Lattal, & Berger, 2007; Flint, et al., 2007; Taubenfeld, et al., 2001; Wu, Lin, Wang, & Hsieh, 2007).

Prior research (with anisomycin, Wanisch & Wotjak, 2008) moreover suggests that subcutaneous administration results in more long-lasting effects than intraperitoneal injection, and may therefore produce inhibitory effects on protein synthesis that are more similar to those of intracerebral drug administration.

### Statistical analyses

Statistical analyses were conducted using Statistica 13.5.0.17 (TIBCO Software Inc., Palo Alto, CA, USA), and the main analyses were confirmed in JASP 0.9.1 (JASP Team). Cohen’s *d* effect sizes were calculated according to (Lakens, 2013) and power (with α = .05) was estimated with G*Power 3.1.9.2 (Faul, Erdfelder, Lang, & Buchner, 2007). Significance levels were set at p < .05. All graphs were created using GraphPad Prism 7.02 (GraphPad Software, La Jolla, CA, USA).

#### Preregistered analyses

In Experiments 1-7, freezing during the Reactivation CS was compared between drug-treated and control groups (two-sided t-tests) to assess whether all animals retrieved the fear memory to a similar extent. Next, the crucial analysis was a comparison of tone fear memory retention on Test 1 (evaluation of freezing during the first 4 CSs in Experiments 1-6 and during the 3 CSs in Experiment 7). Given the clear prediction of the direction of the effect (i.e., drug < control), one-sided t-tests were conducted on the average freezing during these CSs. When Levene’s test suggested a violation of the equal variance assumption, Welch’s t-test was performed (Experiments 4 and 7). To consider all available information from each trial, we also conducted repeated-measures ANOVAs (RM ANOVAs) with factors Trial and Group, if necessary with Greenhouse-Geisser correction, and followed up with Tukey’s post-hoc tests. Additionally, planned analyses regarding acute effects of anisomycin and cycloheximide on preCS and CS-elicited freezing during the short-term memory test (Experiment 7) were performed (two-sided t-tests). As indicated above and in the graphs, most experiments continued after the (first) 3 or 4 CSs of Test 1, with the aim of evaluating the expected amnestic effect in more detail and over a longer period of time. Given the absence of any evidence for amnesia in drug-treated compared to control animals, reporting the results of all additional planned analyses (i.e., comparing the extent of spontaneous recovery, renewal and reinstatement between groups) seems superfluous. For the sake of completeness, all data are, however, shown in the graphs and the results of all preregistered analyses can be found on OSF. In none of these analyses significant group differences were found.

In Experiment 8, the effect on consolidation of fear memory was evaluated by comparing freezing during the first 3 CSs of Test 1 between drug-treated and control rats (RM ANOVA). Rats that had received vehicle after Training (n = 12) were then allocated to two subgroups that received cycloheximide or vehicle after extinction training during Test 1. Extinction of both subgroups during Test 1 (all 12 CSs) was evaluated with a RM ANOVA with Greenhouse-Geisser correction. Extinction memory retention (all 3 CSs) was analyzed with a RM ANOVA.

#### Analyses that were not part of the preregistered analysis plan

Given the results of the preregistered (frequentist inference) analyses of Experiments 1-7, we conducted additional Bayesian analyses, using the BayesFactor package in R (version 3.3.2, R Foundation) and assuming a default Cauchy prior with a scaling factor of .707. In order to quantify the evidence for the absence of a group effect on average freezing during the tones at Test, we performed Bayesian one-sided t-tests (drug < control). In addition to tests per experiment, we carried out a Bayesian fixed-effects, one-sided meta-analysis using the t-values from one-sided t-tests (meta.ttestBF function). For all reported Bayesian analyses, BF_01_ quantifies evidence in favor of the absence of an amnestic effect (i.e., H_0_: drug-treated rats do not show lower freezing than control rats at Test). Bayes factors were categorized in accordance with (Wetzels et al., 2011), with a BF_01_ between 1 and 3 suggesting anecdotal evidence for H_0_, and a BF_01_ between 3 and 10 suggesting substantial evidence.

To more comprehensively characterize the effects of the applied drugs, we also evaluated the acute effects of propranolol on freezing during the Reactivation CS in Experiment 4 (two-sided t-test, no exclusion criterion applied, thus n = 8 per group). A similar analysis was done for acute effects of propranolol on freezing during an extinction session consisting of 12 CSs (RM ANOVA, Luyten et al. in prep). Long-term effects on body weight for rapamycin, anisomycin and cycloheximide (Experiments 5-7) were evaluated by comparing the increase in body weight (difference between last measurement before and after injection) between groups (two-sided t-tests).

For a more thorough analysis of extinction retention in Experiment 8, the change in freezing from the last trial of Test 1 to the first trial of Test 2 was evaluated with a RM ANOVA with factors Test and Group.

#### Power calculations

As mentioned above, published studies using propranolol, anisomycin or cycloheximide typically report large effect sizes. Intraperitoneal injection of 10 mg/kg propranolol after retrieval of a conditioned tone in adult male rats (Debiec & LeDoux, 2004; Muravieva & Alberini, 2010) has repeatedly been found to produce strong amnestic effects with an average Cohen’s *d* = 1.99 (based upon inspection of the graphs of both studies), resulting in a power of .98 when comparing 8 propranolol and 8 control rats (all calculations are for one-sided t-tests, unless indicated otherwise). Power is still >0.90 with only 5 propranolol and 7 control animals, as is the case for Experiment 4.

Protein synthesis inhibitors anisomycin and cycloheximide have both been administered in auditory fear conditioning studies, albeit not systemically, but locally into the basolateral amygdala. They have, however, been applied systemically for example for interference with inhibitory avoidance memories in rats. For anisomycin, the effect size in Nader et al. (2000) and Taubenfeld et al. (2001) (150 mg/kg) was *d* = 1.92 on average. This results in a power of .99 when using a sample size of 10 rats per group. For cycloheximide, the effect size in Duvarci et al. (2005) and Gisquet-Verrier et al. (2015) was *d* = 1.29 on average, resulting in a power of .94 with 14 cycloheximide-treated and 12 control rats.

Effect sizes in rat auditory fear conditioning studies using injections of mTOR inhibitor rapamycin (20-40 mg/kg) are considerably smaller (average *d* = 0.59 in Hoffman, et al., 2015; Y. Li, et al., 2013; Tallot, et al., 2017). The sample sizes used in the current study (n = 8 per group) do not allow for a power of ≥.80 (rather 0.30), but experiments with different doses of rapamycin do permit us to explore whether there is at least a trend supporting an amnestic effect.

In the final study, we investigated the effect of cycloheximide on the consolidation of fear memories. Published reports described significant interference of cycloheximide with the consolidation of tone fear memories (in rats and mice; Kochli, et al., 2015; Lay, et al., 2018; Stiedl, et al., 1999), with *d* = 1.24 on average, yielding a power of 0.83 with 12 animals per group (two-sided t-test). In a subset of animals (n = 6 per group), we evaluated the effect of cycloheximide on the consolidation of an extinction memory. This analysis may be underpowered, although some prior studies did show very large effect sizes (e.g., Ryan, Roy, Pignatelli, Arons, & Tonegawa, 2015), with systemic cycloheximide and contextual fear in mice, *d* = 1.77), which would still yield acceptable power (0.79) with only 6 animals per subgroup (two-sided t-test).

## Results

### No evidence for drug-induced post-retrieval amnesia (‘reconsolidation interference’)

In Experiments 1-4, we aimed to induce post-retrieval amnesia using systemic propranolol (***Fig. 1***), in Experiments 5-6 with rapamycin (***Fig. 2***) and in Experiment 7 with anisomycin or cycloheximide (***Fig. 3***). Relevant statistical analyses are reported in ***Table 1***. In all experiments, drug-treated and control animals showed no significant differences in freezing during the Reactivation CS, suggesting similar retrieval of the cued fear memory in each group. Freezing before the CS during both Reactivation and Test 1 sessions was very low in all experiments, indicative of tone fear memory retrieval that was unconfounded by contextual fear. Cued fear memory retention on Test 1 was evaluated through freezing during the first 3 or 4 CSs. Against our expectations, there was no evidence for amnesia in the drug-treated animals in any of the experiments. Given the lack of any effects of propranolol, rapamycin, anisomycin or cycloheximide, additional Bayesian analyses were carried out that collectively suggested substantial evidence for the absence of an amnestic effect in this series of experiments.

**Figure 2:**
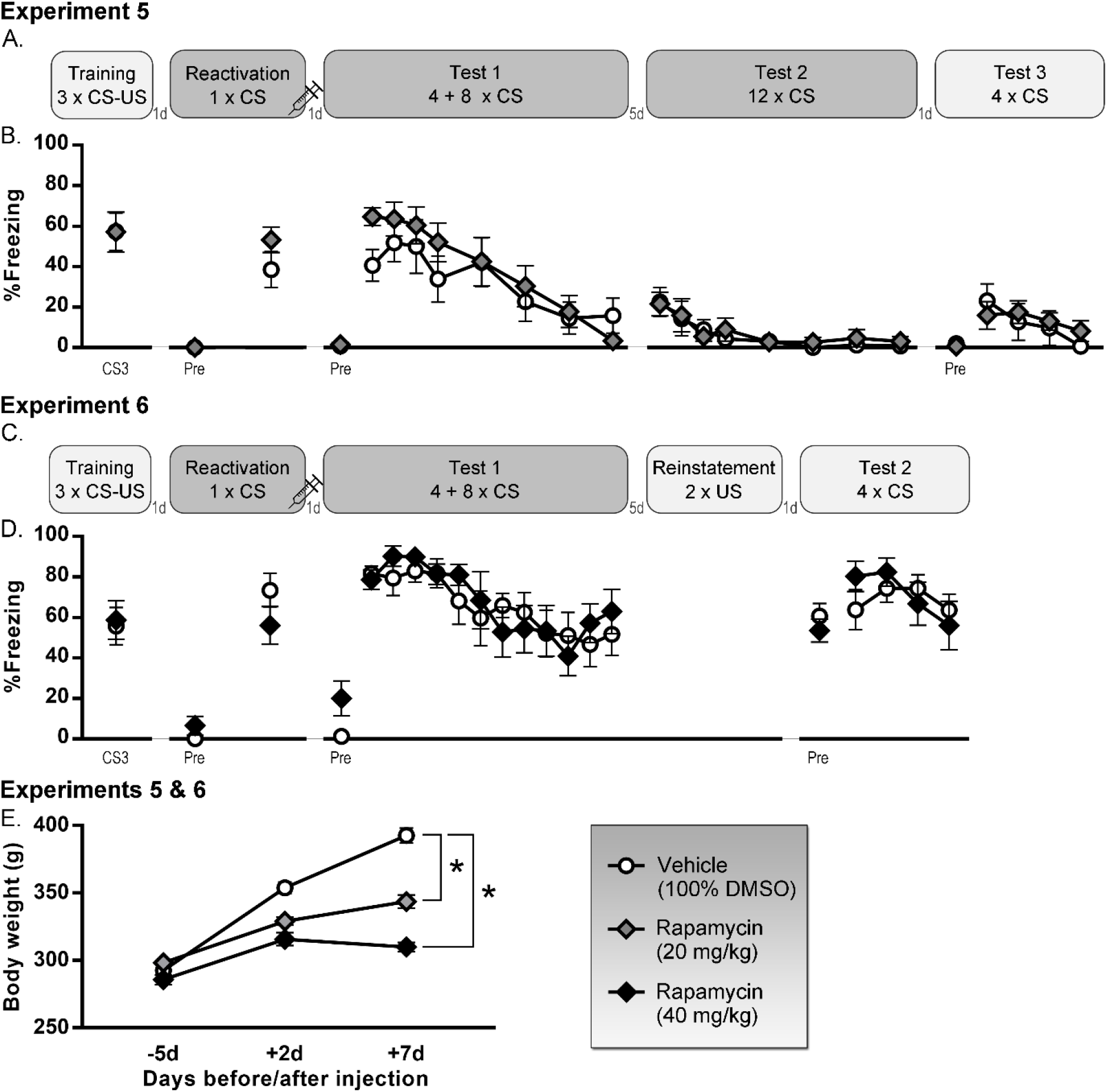
Rapamycin (20-40 mg/kg) experiments. **A-B.** Experiment 5 (20 mg/kg), n = 8 per group. **C-D.** Experiment 6 (40 mg/kg), n = 8 per group. Percentage freezing during each tone (mean and SEM) is shown, except for Test 1 and 2 of Experiment 5, where freezing during CS5-7-9-11 was not measured. ‘Pre’ is contextual freezing before the first tone presentation of the session. **E.** Effects on body weight (mean and SEM) in both experiments, n = 16 vehicle (100% DMSO) rats, n = 8 rapamycin (20 mg/kg) rats and n = 8 rapamycin (40 mg/kg) rats. Light gray box indicates that a session takes place in context A, dark gray is context B. CS: conditioned stimulus, US: unconditioned stimulus, DMSO: dimethyl sulfoxide, d: day(s), *significant group differences (p<.0001).

**Figure 3:**
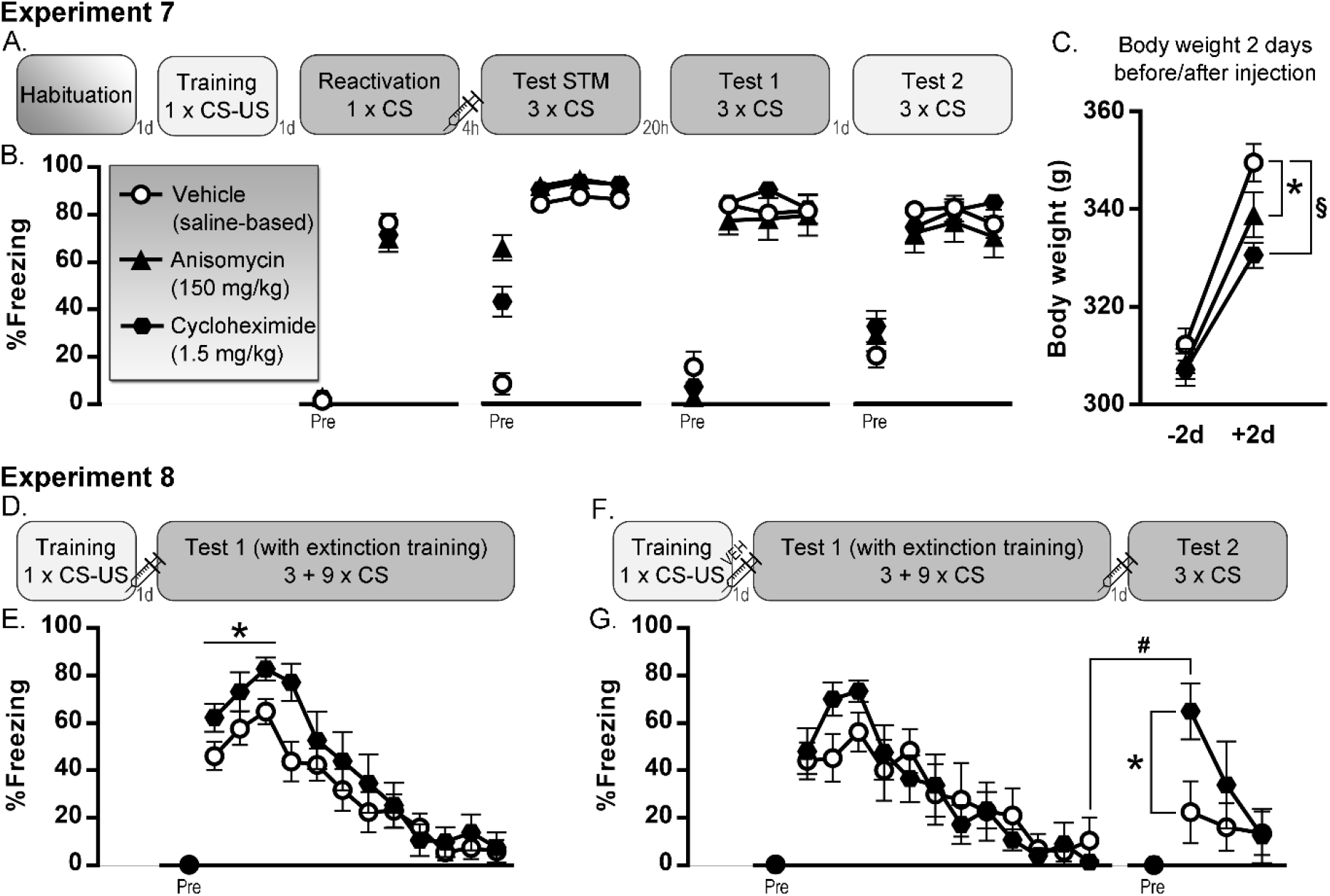
Anisomycin (150 mg/kg) and cycloheximide (1.5 mg/kg) experiments. **A-B.** Experiment 7, n = 12 vehicle (saline-based) rats, n = 10 anisomycin rats and n = 14 cycloheximide rats. **C.** Effects on body weight (mean and SEM) in Experiment 7. **D-E.** Experiment 8, n = 12 per group. **F-G.** Subset of Experiment 8, including animals that received vehicle after training, and subsequently received vehicle (n = 6) or cycloheximide (n = 6) after extinction training. Percentage freezing during each tone (mean and SEM) is shown. ‘Pre’ is contextual freezing before the first tone presentation of the session. Light gray box indicates that a session takes place in context A, dark gray is context B. STM: short-term memory, CS: conditioned stimulus, US: unconditioned stimulus, d: day(s), h: hours, significant group differences (*p<.05, ^§^p<.0001), significant within-group difference (^#^p<.01).

**Table 1:** Statistical evaluation of post-retrieval amnesia with systemic administration of propranolol, rapamycin, anisomycin or cycloheximide provides no evidence in favor of an amnestic effect. Note that BF_01_ quantifies the obtained evidence in favor of the null hypothesis (i.e., absence of an amnestic effect) relative to the alternative hypothesis (i.e., presence of an amnestic effect). For the Bayesian meta-analysis, the comparison of Prop versus Prop NoReact in Experiment 1 was not included as this was a comparison with an additional control condition. Ani: anisomycin (150 mg/kg); Cyclo: cycloheximide (1.5 mg/kg); Prop: propranolol (10 mg/kg); Rap 20: rapamycin (20 mg/kg); Rap 40: rapamycin (40 mg/kg); Sal: saline; Veh: vehicle.

| Experiment | Figure | Drug condition | n | Control condition | n | Reactivation CS (two-sided t-test) | Retention Test | Retention Test (one-sided t-test, drug < control) | Retention Test (RM ANOVA, Trial by Group effect) | Retention Test (Bayesian one-sided t-test, drug < control) | Retention Test (Bayesian evidence against drug < control) | Retention Test (Bayesian one-sided meta-analysis) |
| --- | --- | --- | --- | --- | --- | --- | --- | --- | --- | --- | --- | --- |
| Experiment 1 | 1B | Prop | 8 | Sal | 8 | $t(14) = 0.29$, $p = .78$ | Test 1 (4 CSs) | $t(14) = 0.12$, $p = .55$ | $F(3,42) = 0.96$, $p = .42$ | $BF_{01} = 2.52$ | Anecdotal | $BF_{01} = 9.93$, suggesting **substantial evidence for the absence of an amnestic effect** |
| Experiment 1 | 1B | Prop | 8 | Prop NoReact | 8 | NA | Test 1 (4 CSs) | $t(14) = 0.01$, $p = .50$ | $F(3,42) = 2.02$, $p = .13$ | $BF_{01} = 2.35$ | Anecdotal | |
| Experiment 2 | 1D | Prop | 8 | Sal | 8 | $t(14) = -0.09$, $p = .93$ | Test 1 (4 CSs) | $t(14) = 0.01$, $p = .50$ | $F(3,42) = 0.56$, $p = .64$ | $BF_{01} = 2.35$ | Anecdotal | |
| Experiment 3 | 1F | Prop | 8 | Sal | 6 | $t(12) = -0.13$, $p = .90$ | Test 1 (4 CSs) | $t(12) = 0.84$, $p = .79$ | $F(3,36) = 0.69$, $p = .57$ | $BF_{01} = 3.45$ | Substantial | |
| Experiment 4 | 1H | Prop | 5 | Sal | 7 | $t(10) = -0.37$, $p = .72$ | Test 1 (4 CSs) | $t(4.52) = -0.77$, $p = .24$ | $F(3,30) = 0.24$, $p = .87$ | $BF_{01} = 1.09$ | Anecdotal | |
| Experiment 5 | 2B | Rap 20 | 8 | Veh | 8 | $t(14) = 1.34$, $p = .20$ | Test 1 (4 CSs) | $t(14) = 1.43$, $p = .91$ | $F(3,42) = 0.59$, $p = 0.63$ | $BF_{01} = 4.59$ | Substantial | |
| Experiment 6 | 2D | Rap 40 | 8 | Veh | 8 | $t(14) = -1.38$, $p = .19$ | Test 1 (4 CSs) | $t(14) = 0.51$, $p = .69$ | $F(3,42) = 1.83$, $p = .16$ | $BF_{01} = 3.13$ | Substantial | |
| Experiment 7 | 3B | Ani | 10 | Veh | 12 | $t(20) = -0.97$, $p = .34$ | Test 2 (3 CSs) | $t(10.08) = -0.49$, $p = .32$ | $F(2,40) = 0.33$, $p = .72$ | $BF_{01} = 1.75$ | Anecdotal | |
| Experiment 7 | 3B | Cyclo | 14 | Veh | 12 | $t(24) = -0.91$, $p = .37$ | Test 2 (3 CSs) | $t(24) = 1.04$, $p = .84$ | $F(2,80) = 1.39$, $p = .26$ | $BF_{01} = 4.83$ | Substantial | |

Further extinction, spontaneous recovery, renewal and/or reinstatement of memory expression was evaluated, as illustrated in ***Fig. 1-3***. No group differences were found, as to be expected given the lack of amnestic effects on Test 1.

### Evidence for drug-induced effects on freezing, body weight and memory consolidation

To further characterize the effects of the drugs at the applied dose and administration route in adult male Sprague-Dawley rats, several additional analyses and experiments were carried out. These do suggest that the drugs have detectable effects on freezing behavior, on body weight gain and on consolidation of fear and extinction memories, as detailed below. All relevant statistical analyses are reported in ***Table 2***.

**Table 2:**
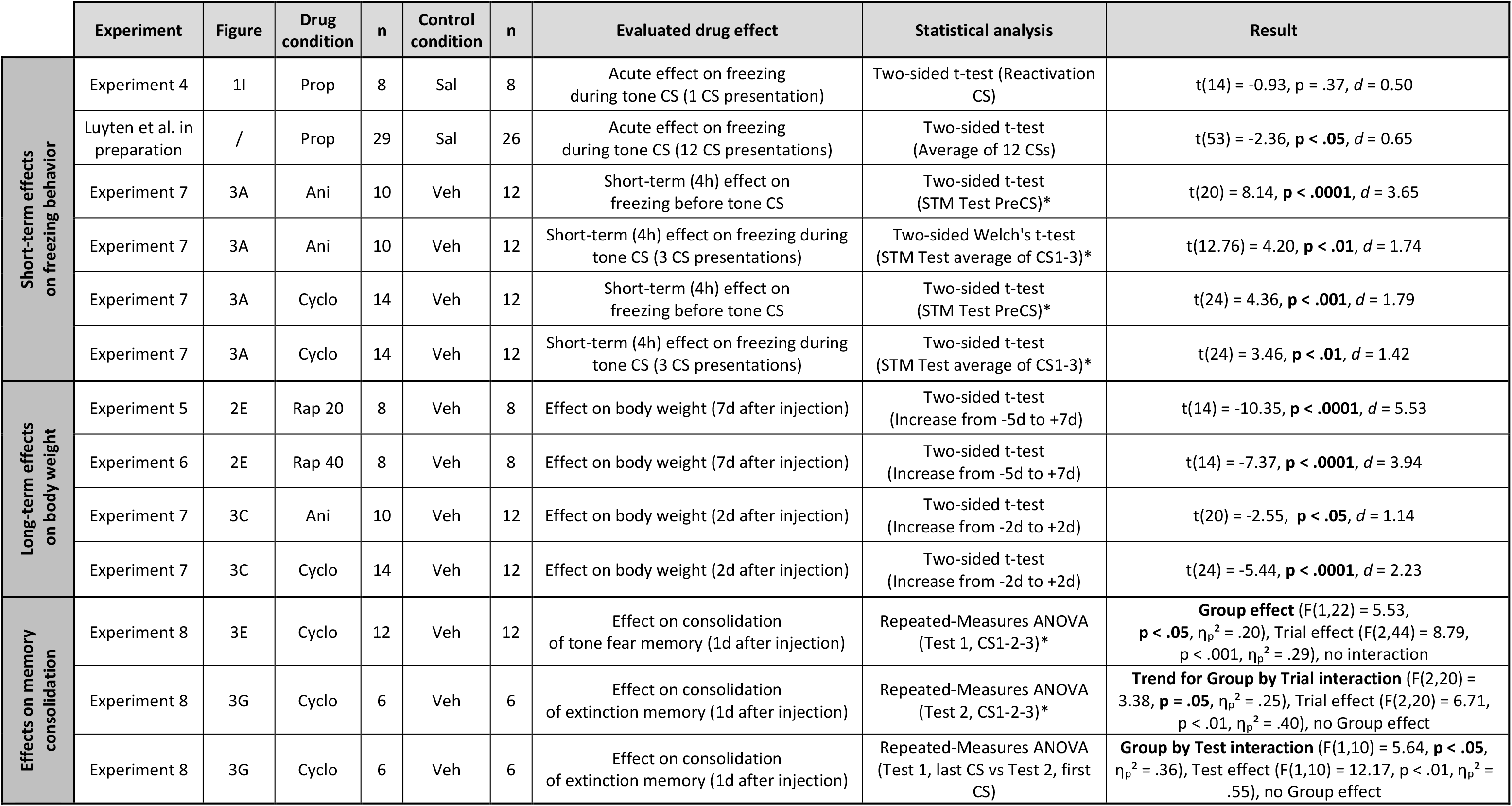
Statistical evaluation of other effects of propranolol, rapamycin, anisomycin or cycloheximide suggests that the drugs (same dose and route as in Table 1) do have detectable physiological effects. Ani: anisomycin (150 mg/kg); Cyclo: cycloheximide (1.5 mg/kg); Prop: propranolol (10 mg/kg); Rap 20: rapamycin (20 mg/kg); Rap 40: rapamycin (40 mg/kg); Sal: saline; Veh: vehicle; *preregistered analysis.

#### Acute drug effects on freezing behavior

In Experiment 4, propranolol was administered prior to memory reactivation, allowing us to evaluate its effects on freezing during the CS (***Fig. 1I***). Freezing in the Prop group was lower than in the control condition, but the effect was too small to be statistically significant with a sample size of 8 per group. In a recent study, we evaluated the effect of propranolol injection 20 minutes before a session with 12 CS presentations in 29 Prop and 26 Sal rats, and did find significantly lower freezing in the Prop group (Luyten et al., in preparation). In addition, we found effects of anisomycin and cycloheximide during the short-term memory test 4 hours after injection. The analyses are shown in ***Table 2*** and expounded in the ***Supplement***.

#### Long-term effects of rapamycin, anisomycin and cycloheximide on body weight

Interference with protein synthesis on a systemic level has certain toxic effects (see also ***Supplement***), and although not lethal at the applied dose, effects on body weight were anticipated. Indeed, when comparing the increase in body weight from the last measurement before until the last measurement after injection, animals that had received rapamycin (***Fig. 2E***), anisomycin or cycloheximide (***Fig. 3C***) gained significantly less weight than control animals.

#### Effects of cycloheximide on consolidation of fear and extinction memories

In Experiment 8, we assessed the effects of cycloheximide on consolidation rather than reconsolidation. We hypothesized that injection of cycloheximide after fear acquisition would interfere with memory consolidation, resulting in decreased freezing during the first 3 CSs on Test 1 (***Fig. 3E***). To our surprise, we found the opposite effect, with more freezing in the cycloheximide rats. It is unlikely that this was a non-specific effect of cycloheximide on freezing, given that no such increases were observed during the period preceding the first Test 1 CS in Experiment 8, nor on Test 1 in Experiment 7 (which took place 24h after injection). A subset of rats was further tested in Experiment 8 to assess the effects of cycloheximide on extinction memory consolidation (***Fig. 3G***). During extinction (full Test 1 session) both subgroups were equivalent and extinguished to the same degree, after which they received an injection with cycloheximide or vehicle. Our hypothesis that cycloheximide would interfere with consolidation of the extinction memory and thus result in more freezing on Test 2 was confirmed and particularly evident on the first trial of this session, where vehicle animals showed good retention of the extinction memory, whereas cycloheximide animals did not.

## Discussion

The aim of this series of experiments was to establish a protocol that could be used to probe the mechanisms of post-retrieval amnesia. We thus attempted to conceptually and exactly replicate prior published studies in which administration of propranolol, rapamycin, anisomycin or cycloheximide upon auditory fear memory retrieval resulted in a significant attenuation of the fear response during subsequent tests (i.e., drug-induced post-retrieval amnesia). We adhered to universal behavioral procedures and pharmacological agents, including a direct replication attempt of Debiec & LeDoux (2004), using sample sizes that were expected to yield sufficient power. In light of the current literature (e.g., Debiec & LeDoux, 2004; Haubrich et al., 2015; Mac Callum, et al., 2014; Nader, et al., 2000), which contains an abundance of studies that found post-retrieval amnesia, often with large effect sizes, and hardly any failures to replicate under standard training and reactivation conditions (one exception is Pitman, et al. (2001) whose graph suggested no amnestic effect of propranolol but did not report any formal statistical analyses supporting this failure). Therefore, we anticipated to readily reproduce the effect in our laboratory. Against our expectations, we did not find any evidence for an amnestic effect, and Bayesian analyses even indicated substantial evidence for the absence of an effect (***Table 1***). Inasmuch as we administered all drugs systemically and confined this endeavor to auditory fear memories, our conclusions cannot surpass these procedural choices. Nevertheless, in a recent study we also described repeated failures to find post-retrieval amnesia in contextual fear conditioning studies with Wistar and Sprague-Dawley rats, using either propranolol or midazolam injections (Schroyens, Alfei, Schnell, Luyten, & Beckers, 2019), suggesting that these reproducibility problems may extend to other fear conditioning procedures and pharmacological agents as well.

The lack of amnestic effects in the current paper may have several causes (Reichelt & Lee, 2013). Drug-induced post-retrieval amnesia entails two crucial elements: a suitable drug and appropriate retrieval. First, it is possible that the pharmacological agents that we administered were all ineffective in interfering with memory retention. Second, the various behavioral procedures that we used might not have resulted in actual memory reactivation, understood as a destabilization of the memory trace which is required in order to interfere with it, as proposed by the reconsolidation blockade hypothesis (Nader, et al., 2000). Instead of having this intended effect, our reactivation session may have led to mere retrieval, without destabilization (Sevenster, Beckers, & Kindt, 2012). We cannot be absolutely sure if one or both factors account for our results, and there are, of course, countless parametric variations that we did not try. Nevertheless, below we will argue why such ‘easy’ explanations for our data and conclusions are debatable.

First, in order to evaluate whether the drugs had any biological effects at the applied dose and administration route, we performed several additional tests and analyses (***Table 2***). We observed acute effects of propranolol, anisomycin and cycloheximide on freezing behavior. The effects of both protein synthesis inhibitors most likely reflect transient and non-specific changes in general mobility of the animals, in line with signs of sickness that appear shortly after injection of such drugs (***Supplement***; Blaiss & Janak, 2007; Gisquet-Verrier et al., 2015). Decreased freezing following propranolol administration presumably does represent a genuine reduction in fear expression (Rodriguez-Romaguera, Sotres-Bayon, Mueller, & Quirk, 2009), and is in line with its beta-blocking effects which produce an attenuation of heart rate and blood pressure (Chalkia, Weermeijer, Van Oudenhove, & Beckers, 2019). In addition to these effects on freezing behavior, we found that the protein synthesis inhibitors (i.e., rapamycin, anisomycin and cycloheximide) had long-lasting effects on body weight. After a single injection, animals in the drug conditions gained significantly less weight than vehicle controls over a period of several days, in line with prior studies (Adamec, Strasser, Blundell, Burton, & McKay, 2006; Hebert et al., 2014). Last but not least, we investigated whether we were able to interfere with consolidation, rather than reconsolidation (Experiment 8). We used cycloheximide in this study because of its lower toxicity compared to anisomycin (***Supplement***), the anticipated larger effects in comparison with rapamycin (see effect sizes above) and the documented ineffectiveness of propranolol to interfere with consolidation of auditory fear (Debiec & LeDoux, 2004). We first examined the effect on consolidation of fear memory, and found an unexpected but significant effect in the opposite direction, i.e., better retention (indexed as higher freezing) at the start of the test session in drug-treated animals. Importantly, when we next investigated the effect of cycloheximide on consolidation of the extinction memory, we did indeed find evidence for amnesia. Animals that had received the protein synthesis inhibitor showed worse retention than controls, indicative of interference with the consolidation of fear extinction. All in all, we found clear short- and long-term behavioral and physiological effects of the pharmacological agents that we used, along with effects on consolidation, arguing against the possibility that our drugs were generally ineffective using the administration route and dose that we applied in Experiments 1-7.

Second, as mentioned above, we cannot exclude that, in Experiments 1-7, we failed to induce the destabilization that is thought to be required in order for the retrieved fear memory to undergo changes that can result in amnesia. Although our memories were presumably not too old or too strong (two conditions that may hamper post-retrieval amnesia (Suzuki, et al., 2004; Wang, et al., 2009)), and our animals clearly retrieved the fear memory during the reactivation session (moderate to high freezing upon presentation of the conditioned tone), we cannot be sure that the memory destabilized. Prior research supports that such destabilization only occurs when the reactivation session involves an appropriate degree of prediction error, which is a function of the extent to which the contingencies at the time of retrieval match the contingencies at the time of memory acquisition (Schultz & Dickinson, 2000). We can assume that, in our experiments, the mismatch between the unreinforced reactivation tone and the preceding acquisition session (100% reinforced tones) was large enough to create the amount of prediction error that appears to be required for memory destabilization and subsequent interference with its restabilization (Pedreira, Perez-Cuesta, & Maldonado, 2004; Sevenster, Beckers, & Kindt, 2013). Moreover, given the similarity of our protocols and the observed freezing levels during the reactivation session (see ***Supplement***) with those of published reports, it is unlikely that there was no such mismatch.

A noteworthy difference between previous publications and our work is that the animals were obtained from a different supplier. Our rats were of the same strain, age and sex as in many prior reports, but we cannot exclude that small genetic variations (Theilmann et al., 2016) or subtle differences in early-life experiences (Schroyens, Bender, et al., 2019) influenced susceptibility to drug-induced post-retrieval amnesia. Laboratories that reliably find amnestic effects could examine this in more detail, by comparing post-retrieval amnesia in (different strains of) rodents from different suppliers, or by investigating the effects of genetic or environmental manipulations on the success of this intervention. If such small (epi)genetic variations turn out to be crucial, this observation has far-reaching implications for the overall generalizability of drug-induced post-retrieval amnesia of fear memories. Our replication failures indeed suggest that obtaining amnestic effects depends on very subtle differences between experiments. This makes the current lack of published null findings even more unfortunate, because the field needs information about the conditions under which the effect is not found as much as the success stories, in order to identify these unknown conditions that may influence memory malleability.

As mentioned above, the conclusions of this paper are limited to systemic drug administration and cued fear memory, a deliberate choice in light of the direct relevance of such procedures to clinical applications, e.g., for traumatized patients (Kindt & Van Emmerik, 2016). It is absolutely possible that local drug infusion (e.g., in the amygdala, cf Duvarci et al. (2005) and Nader et al. (2000)) would have permitted us to observe amnestic effects, given that intracerebral administration – albeit clinically less relevant for the time being – provides a temporal and spatial precision that cannot be achieved with systemic administration. In addition, it is possible that the cannulated rats that were tested in part of the prior literature behaved or even learned somewhat differently compared to our non-operated animals. Bear in mind however that the large majority of the studies that we present here rely on protocols that have been used often and successfully for the induction of post-retrieval amnesia using systemic administration in non-operated rats.

An additional caveat is that, even though our experiments with propranolol, anisomycin and cycloheximide were well-powered in view of the effects reported in the literature, those effect sizes may be a gross overestimation of the true effect (Button et al., 2013). Even a true effect will occasionally lead to a non-significant finding, given the nature of the statistical analyses that are typically used to support the presence of the effect (α = .05). That being said, it is implausible that such statistical considerations may comprehensively account for the lack of amnesia that we observed here. We found not even a trend for amnesia and, moreover, Bayesian analyses suggested substantial evidence for the absence of such effect.

Finally, we note that attempts to pharmacologically induce post-retrieval amnesia in other aversive learning procedures, such as inhibitory avoidance or immediate shock conditioning after context pre-exposure, have met with varying degrees of success (Biedenkapp & Rudy, 2004; Muravieva & Alberini, 2010; Reichelt & Lee, 2013). Our results also fit with the mixed findings in the human cued fear conditioning literature. Although several human fear conditioning studies have found propranolol-induced post-retrieval amnesia (Kindt, et al., 2009; Schwabe, Nader, Wolf, Beaudry, & Pruessner, 2012), others reported failures to replicate this result (Bos, et al., 2014; Chalkia, et al., 2019; Thome et al., 2016). Furthermore, our series of null findings in rodents aligns with personal communication from several researchers in the field and reports in unpublished dissertations (e.g., Vousden, 2017) of failures to find drug-induced post-retrieval amnesia in rat or mouse fear conditioning under standard training and reactivation conditions. This striking contrast with most of the existing rodent literature suggests a reporting bias, although we cannot substantiate such claim with the current study alone. Moreover, we do not intend to question the veracity of this phenomenon as a whole. Rather, our data should be seen as an important warning against overly enthusiastic statements regarding the generalizability and clinical translatability of drug-induced post-retrieval amnesia.

## Acknowledgements

We thank Dr. Kelly Luyck for her technical assistance with the preparation of the anisomycin and cycloheximide solutions. We acknowledge the financial support of the European Research Council (ERC Consolidator Grant to T. Beckers, grant number 648176).

## Supplement

### 1. Details regarding behavioral procedures

All procedural details can be found in the preregistrations on the Open Science Framework (OSF) (https://osf.io/j5dgx). The most important settings and parameters are also listed below.

#### 1.1 Conditioned stimulus (CS) and unconditioned stimulus (US) characteristics

**Table S1:**
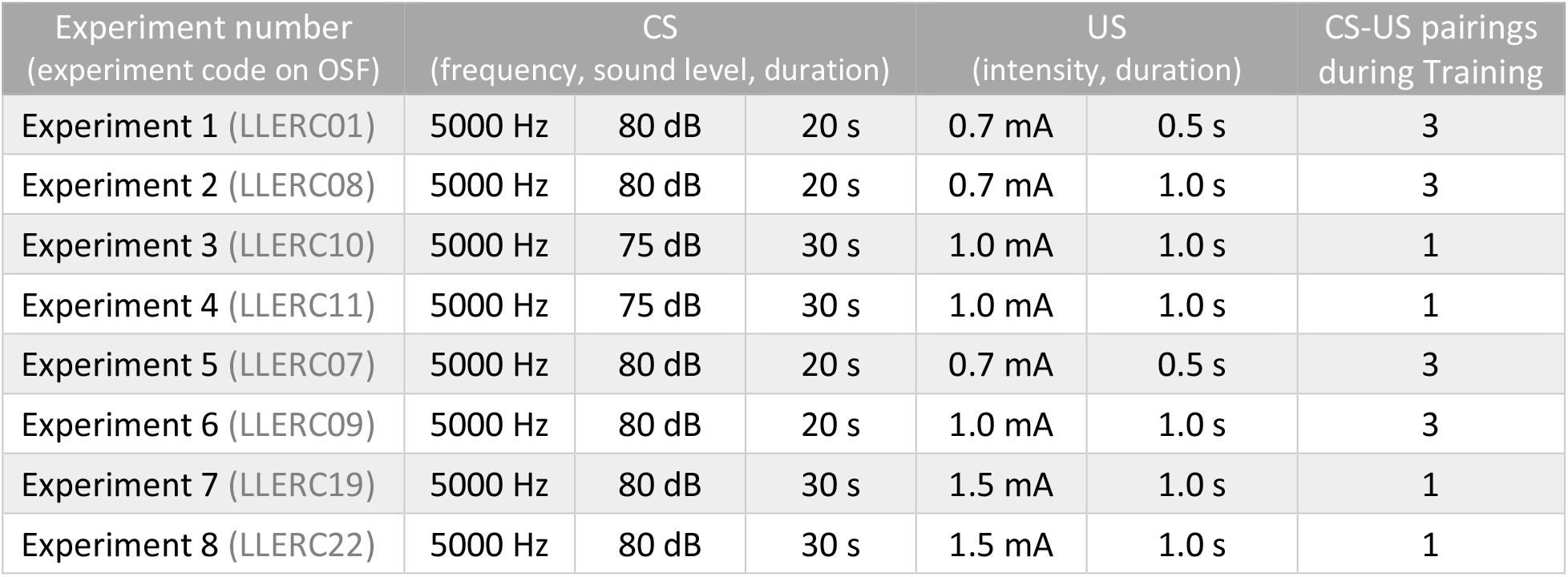
Parameters of conditioned (CS) and unconditioned stimuli (US) in Experiments 1-8.

#### 1.2 Timing of stimulus presentations during behavioral test sessions

##### Experiment 1

**Habituation**: 600 s context exposure

**Training**: 120 s acclimation, interval between CS onsets is 180 s, rat removed from context 120 s after last CS onset

**Reactivation**: 120 s acclimation, rat removed from context 60 s after CS onset

**Test 1-2-3**: 120 s acclimation, interval between CS onsets is 120 s, rat removed from context 120 s after last CS onset

##### Experiment 2

**Training**: 600 s acclimation, interval between CS onsets is 180 s, rat removed from context 180 s after last CS onset

**Reactivation**: 120 s acclimation, rat removed from context 60 s after CS onset

**Test 1-2**: 120 s acclimation, interval between CS onsets is on average 120 s (range 100-140 s), rat removed from context 120 s after last CS onset

**Reinstatement**: 300 s acclimation, interval between US onsets is 180 s, rat removed from context 120 s after last US onset

##### Experiment 3

**Habituation**: 600 s context exposure

**Training**: 120 s acclimation, 1 CS-US pairing, rat removed from context 60 s after CS onset

**Reactivation**: 120 s acclimation, 1 CS, rat removed from context 60 s after CS onset

**Test 1-2-3**: 120 s acclimation, 4 CSs and interval between CS onsets is on average 120 s (range 90-150 s), rat removed from context 60 s after last CS onset

##### Experiment 4

**Training**: 480 s acclimation, 1 CS-US pairing, rat removed from context 60 s after CS onset

**Reactivation**: 120 s acclimation, 1 CS, rat removed from context 60 s after CS onset

**Test 1**: 120 s acclimation, 4 CSs and interval between CS onsets is on average 120 s (range 90-150 s), rat removed from context 60 s after last CS onset

##### Experiment 5

**Training**: 600 s acclimation, interval between CS onsets is 180 s, rat removed from context 180 s after last CS onset

**Reactivation**: 120 s acclimation, rat removed from context 60 s after CS onset

**Test 1-2-3**: 120 s acclimation, interval between CS onsets is on average 120 s (range 100-140 s), rat removed from context 120 s after last CS onset

##### Experiment 6

**Training**: 600 s acclimation, interval between CS onsets is 180 s, rat removed from context 180 s after last CS onset

**Reactivation**: 120 s acclimation, rat removed from context 60 s after CS onset

**Test 1-2**: 120 s acclimation, interval between CS onsets is on average 120 s (range 100-140 s), rat removed from context 120 s after last CS onset

**Reinstatement:** 300 s acclimation, interval between US onsets is 180 s, rat removed from context 120 s after last US onset

##### Experiment 7

**Habituation**: 600 s context exposure (4-hour interval between two contexts, counterbalanced order)

**Training**: 300 s acclimation, rat removed from context 60 s after CS-US offset

**Reactivation**: 300 s acclimation, rat removed from context 60 s after CS offset

**Test STM-1-2**: 300 s acclimation, interval between CS onsets is 90 s, rat removed from context 60 s after last CS offset

##### Experiment 8

**Training**: 300 s acclimation, rat removed from context 60 s after CS-US offset

**Test 1-2**: 300 s acclimation, interval between CS onsets is 90 s, rat removed from context 60 s after last CS offset

### 2. Experiment S7: Selection of the behavioral procedure for Experiment 7

In Experiment 7, we aimed to evaluate the amnestic effects of two frequently used protein synthesis inhibitors: anisomycin and cycloheximide. Although often employed to interfere with fear memories, we did not find published accounts of systemic administration of these drugs in a cued fear conditioning procedure in rats. As mentioned in the main text, both drugs have been applied systemically in several reports, and have been shown to produce amnesia in various behavioral procedures (e.g., Bernardi et al. 2007; Flint et al. 2007; Haubrich et al. 2015; Milekic & Alberini 2002; Taubenfeld et al. 2001; Wu et al. 2007), but have not yet been investigated in cued fear conditioning. In order to increase our chances of finding an amnestic effect, we therefore looked for papers with a cued fear conditioning procedure and local (intra-amygdala) infusion of these drugs, and conducted a dry run with these procedures (i.e., Experiment S7, vehicle animals only), with the purpose of selecting a behavioral protocol with similar freezing behavior in our hands as in the publications.

In a group of 8 animals, we aimed to replicate the procedure used by Duvarci et al. (2005, Fig. 1) as well as Nader et al. (2000, Fig. 5), a protocol that allowed them to induce post-retrieval amnesia with cycloheximide and anisomycin infusion, respectively. In what follows, this procedure is referred to as the Duvarci 2005 protocol. In a concurrently tested set of 8 other animals, we used the procedure of Nader et al. (2000, Fig. 2c, referred to as the Nader 2000 protocol), which also provided evidence for post-retrieval amnesia after anisomycin infusion, but did not feature a short-term memory test in between the Reactivation session and the long-term memory retention test (Test 1). We preferred a behavioral procedure that did include such a short-term memory (STM) test, because it would provide more information about the nature of the drug effects, if any.

Note that the vehicle used in this supplementary Experiment S7 was 60% DMSO (3 ml/kg DMSO and 2 ml/kg saline), which we intended to use in Experiment 7, but ultimately could not, because anisomycin did not dissolve, in contrast with Sigma-Aldrich’s product information.

Timing of the sessions and stimuli (***Table S2***) was the same as what was eventually used in Experiment 7, except that the exact replication of the Nader 2000 protocol in Experiment S7 did not include a Habituation phase nor Test STM.

**Table S2:** Parameters of conditioned (CS) and unconditioned stimuli (US) in Experiment S7.

| Experiment number<br>(experiment code on OSF) | CS<br>(frequency, sound level, duration) |  |  | US<br>(intensity, duration) |  | CS-US pairings<br>during Training |
| --- | --- | --- | --- | --- | --- | --- |
| Experiment S7<br>(Duvarci 2005 protocol)<br>(LLERC16) | 5000 Hz | 80 dB | 30 s | 1.5 mA | 1.0 s | 1 |
| Experiment S7<br>(Nader 2000 protocol)<br>(LLERC16) | 5000 Hz | 75 dB | 30 s | 2.0 mA | 1.0 s | 1 |

Both behavioral procedures (***Fig. S1***) seemed to replicate quite well in our hands, and rats showed relatively limited between-subject variability in freezing. Given the added value of a short-term memory test, where we expected no amnestic effects of the drugs, we selected the Duvarci 2005 protocol (shown in the top panels of ***Fig. S1***) to be used in Experiment 7.

Mean freezing levels during the Reactivation CS, Test STM (average of CS1-3) and Test 1 (average of CS1-3) were overall similar, but somewhat higher in our hands (Experiment S7, Duvarci 2005 protocol: 56%, 75% and 71%, respectively) than in the control animals of prior publications (Nader et al. 2000: 66%, 54% and 56%, Duvarci et al. 2005: 43%, 44% and 44%, respectively, values obtained from graphs). Such generally higher freezing levels in non-operated animals are not uncommon, although it is not clear whether they reflect differences in fear or differences in freezing tendencies (Luyten et al. 2016; Zhang et al. 2001). Regardless, our data do indicate clear retrieval of the fear memory, no extinction resulting from the short-term memory test and sufficient room to observe amnesia in Experiment 7.

**Figure S1:**
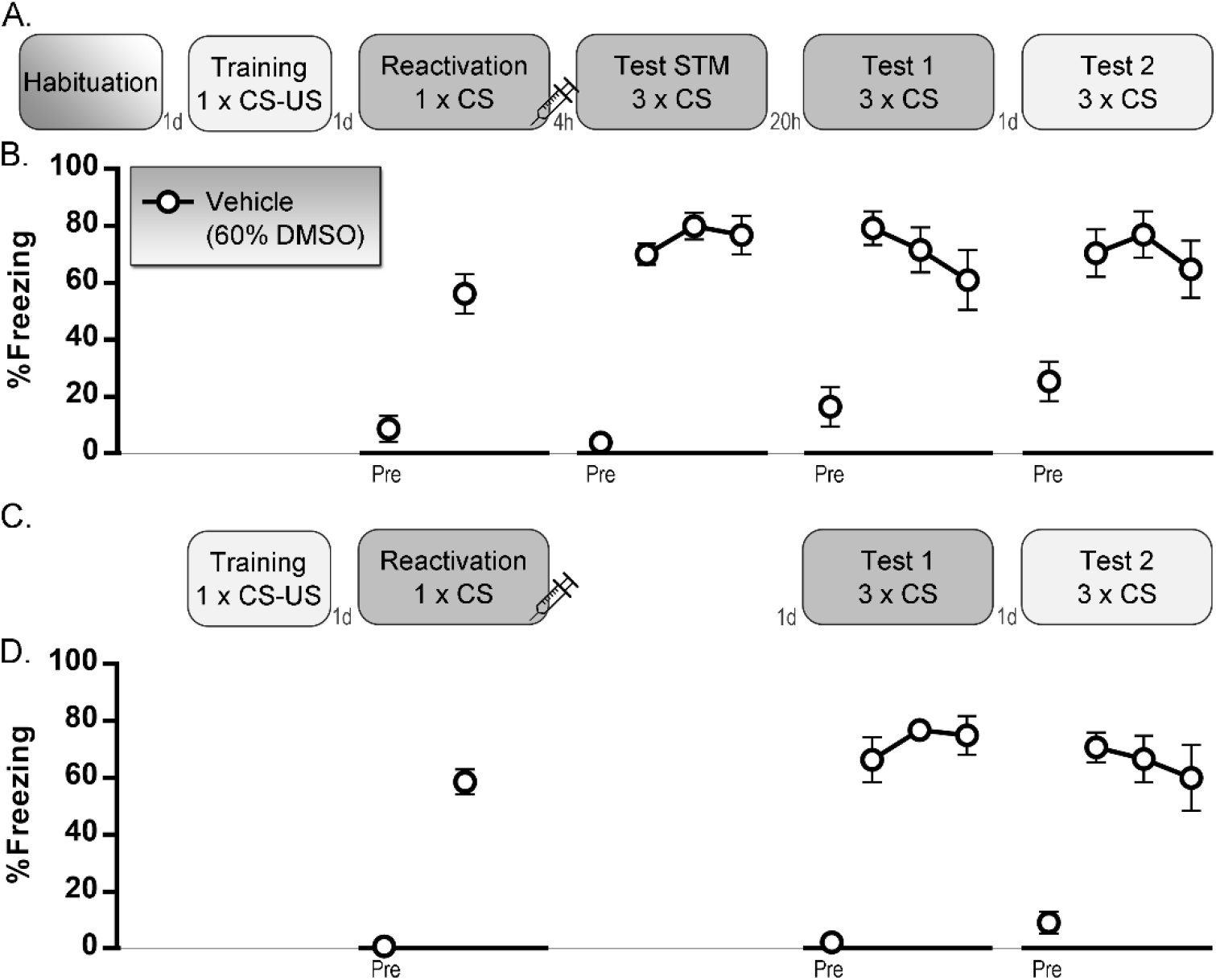
Experiment S7. **A-B.** Duvarci 2005 protocol, n = 8 vehicle rats. **C-D.** Nader 2000 protocol, n = 8 vehicle rats. Percentage freezing during each tone (mean and SEM) is shown. ‘Pre’ is contextual freezing before the first tone presentation of the session. Light gray box indicates that a session takes place in context A, dark gray is context B. STM: short-term memory, CS: conditioned stimulus, US: unconditioned stimulus, DMSO: dimethyl sulfoxide, d: day, h: hours.

### 3. Adverse reactions to the applied drugs

As shown in the main article (***Fig. 2E*** and ***Fig. 3C***), rapamycin, anisomycin and cycloheximide had long-term effects on body weight, with lower weights in drug-treated animals than controls several days after injection.

Furthermore, we observed an increase in freezing(-like) behavior during the short-term memory test in Experiment 7 (***Fig. 3B*** and ***Table 2***). Freezing was scored by an observer blinded to group allocation, but it seems that acute (i.e., 4 hours after injection) side effects may have confounded the freezing measurements during this particular test (see below). Whether these effects have a sedative, numbing, nauseous or other nature is impossible to say, but we can assume that, to a certain degree, the resulting immobility was mistakenly identified as freezing. This is especially clear during the PreCS measurement on Test STM (***Fig. 3B***), where vehicle animals show very little freezing (8% on average), whereas animals that had received a protein synthesis inhibitor 4 hours earlier, were classified as more immobile, anisomycin animals (66%) even to a greater extent than cycloheximide animals (43%). During the CS presentations, the differences are much smaller, presumably due to a ceiling effect (high freezing in vehicle animals as well). Do note that these effects had disappeared by the next day (PreCS freezing on Test 1), suggesting that the animals had recovered sufficiently and that the disruptive acute effects that were seen a few hours after injection were no longer relevant 20 hours later, implying a reliable behavioral read-out for all animals on Test 1 and 2. This was confirmed by the absence of effects on contextual freezing in Test 1 and Test 2 of Experiment 8 as well (***Fig. 3E, G***).

In addition to analyzing the acute effects that were just described, we carried out a qualitative assessment of the animals’ wellbeing in the hours and days after injection, given the potential toxic effects of systemic drugs that are intended to interfere with protein synthesis. Manufacturers’ safety sheets (based on the Registry of Toxic Effects of Chemical Substances) and the literature provided the following toxicity information: LD_50_ DMSO (intraperitoneal, rat): 9.9 ml/kg (Bartsch et al. 1976), LD_50_ rapamycin (intraperitoneal, rat): 18.2 mg/kg; LD_50_ anisomycin (subcutaneous, rat): 230 mg/kg; LD_50_ cycloheximide (subcutaneous, rat): 2.5 mg/kg. Note that the supposed LD_50_ of rapamycin is rather unexpected, given our own and others’ experiences with this drug at doses of 20-40 mg/kg (Hoffman et al. 2015; Li et al. 2013; Tallot et al. 2017). Adverse reactions to systemic administration of the protein synthesis inhibitors have only rarely been reported in prior publications. A few did mention that 150 mg/kg anisomycin produced signs of distress (e.g., lethargy, balance problems, piloerection, hunched back posture) in the hours after injection and weight loss in the longer run (Blaiss & Janak, 2007; Hernandez & Kelley, 2004). Gisquet-Verrier et al. (2015) reported that two rats died after an intraperitoneal injection with 2.8 mg/kg cycloheximide (LD_50_: 3.7 mg/kg).

In our own studies, we made qualitative assessments that were not embedded in the preregistered experimental design, but which were all observations made in the course of the experiment, by a researcher blinded to group division.

**Rapamycin** at a dose of 20 mg/kg (Experiment 5) did not seem to elicit important adverse reactions, apart from the above-mentioned reduced body weight that was seen with both applied doses. In Experiment 6, where we used 40 mg/kg rapamycin, 4 out of 8 drug-treated animals presented with light and two with more pronounced diarrhea one day after injection. Note that, in addition, two out of 8 vehicle animals had light diarrhea, suggesting that the gastrointestinal effects may be ascribed mainly to rapamycin, but partly also to the DMSO vehicle. On the following days, no adverse reactions were noted.

In Experiment 7, all animals were responsive and showed normal muscular tonus when taken out the home cage for the short-term memory test, 4 hours after injection. Nevertheless, careful observation of the animals in the hours after injection did indicate slight to pronounced orbital tightening in 9 out of 10 **anisomycin**-treated rats. Eight of these also showed diarrhea, and two of them were even briefly lying on their back when checked about one hour after injection. Note that all animals seemed to have recovered sufficiently by the next day, and no behavioral abnormalities, nor signs of pain or distress were observed on the day of Test 1. Nine anisomycin animals did show slight skin irritation at the site of injection when examined after euthanasia. Three out of 14 **cycloheximide**-treated rats showed orbital tightening in the hours after injection, and one of these animals also had diarrhea. All animals seemed to have recovered sufficiently by the next day (Test 1 session). Finally, as described above, both anisomycin and cycloheximide injections resulted in lower body weight gain.

### 4. Supplementary discussion regarding indices of destabilization

As mentioned in the main text, we cannot exclude that we failed to destabilize the tone fear memory, which is thought to be a requirement for inducing amnesia. The only possible way to compare our behavioral procedures with those of the publications that they were based upon, is to look at the freezing levels, as an indication of how well we reproduced the behavior in the control condition. In Experiment 3, which was an exact replication attempt of (Debiec & LeDoux, 2004), mean freezing levels during the Reactivation CS and Test 1 (average of CS1-4) in the control condition were very similar in our study (67% and 57%, respectively) and theirs (75% and 67%, respectively, values obtained from graphs), suggesting that the protocol produced comparable levels of fear in our hands. Similar observations can be made for Experiment 7, where the behavioral procedure was an exact replication of Duvarci et al. (2005) and Nader et al. (2000). In other words, our exact replications, and the same holds for our other experiments, showed freezing levels that were comparable to those of successful studies in the literature, suggesting clear retrieval of the fear memory and sufficient room to observe amnesia at test.

Unfortunately, freezing may not suffice as an index of reactivation (Ben Mamou et al. 2006). Given the limitations of behavioral read-outs, prior research has looked for biomarkers that may indicate whether a memory is susceptible to post-retrieval modification. Although such markers currently have little translational value, they may offer insights into what is going on in our animals’ brains and may even help to optimize procedural parameters for destabilization. Some have tried to estimate the probability that the memory will undergo destabilization, e.g., by quantifying NR2B NMDA-receptor subunits in the basal and lateral amygdala after training (Wang et al. 2009). Others have looked at plasticity during the reactivation session and found changes in the expression of the immediate early gene zif-268 in the lateral amygdala (Diaz-Mataix et al. 2013) or in the amount of polyubiquitination (a marker of protein degradation) in the entire amygdala (Jarome et al. 2011). However interesting, in order for such biomarkers to be informative, one in principle requires a comparison group that does show post-retrieval amnesia. Given that this was never the case in our hands, biomarker detection would not have been helpful in delineating adequate reactivation conditions. Moreover, we should not overestimate the decisive value of such markers and keep in mind that they are likely dependent on the behavioral procedure that is being used, the brain region under investigation and perhaps even prior experience of the subject (Finnie & Nader, 2012; Reichelt & Lee, 2013).

